# One for all – Human kidney Caki-1 cells are highly susceptible to infection with corona- and other respiratory viruses

**DOI:** 10.1101/2022.10.25.513760

**Authors:** A. Daniels, S. Fletcher, H. Kerr, A. Kratzel, R.M. Pinto, N. Kriplani, N. Craig, C.J. Hastie, P. Davies, P. Digard, V. Thiel, C. Tait-Burkard

## Abstract

*In vitro* investigations of host-pathogen interactions of viruses are reliant on suitable cell and tissue culture models. Results are only as good as the model they have been generated in. However, choosing cell models for *in vitro* work often depends on what is available in labs or what has previously been used. Despite the vast increase in coronavirus research activity over the past few years, researchers are still heavily reliant on: non-human cells, for example Vero E6, highly heterogeneous or not fully differentiated cells, such as Calu-3, or naturally unsusceptible cells requiring overexpression of receptors and other accessory factors, such as ACE2 and TMPRSS2. Complex cell models, such as primary cell-derived air-liquid interface epithelial models are highly representative of human tissues but are expensive and time-consuming to develop and maintain. They have limited suitability for large-scale and high-throughput analysis.

Using tissue-specific expression pattern as a selection criteria, we identified human kidney cells as an ideal target for severe acute respiratory syndrome coronavirus-2 (SARS-CoV-2) and broader coronavirus infection. We show the use of the highly characterized human kidney cell line Caki-1 for infection with three human coronaviruses: *Betacoronaviruses* SARS-CoV-2 and Middle Eastern respiratory syndrome coronavirus (MERS-CoV) and *Alphacoronavirus* human coronavirus 229E (hCoV-229E). Caki-1 cells show equal or superior susceptibility to all three coronaviruses when compared to other commonly used cell lines for the cultivation of the respective virus. Antibody staining against SARS-CoV-2 N protein shows comparable replication rates. Using a panel of 21 antibodies in infected Caki-1 cells using immunocytochemistry shows the location of viral proteins during replication. In addition, Caki-1 cells were found to be susceptible to two other human respiratory viruses, influenza A virus and respiratory syncytial virus, making them an ideal model for cross-comparison of not only a broad range of coronaviruses but respiratory viruses in general.

**Author Summary:** Investigating how viruses interact with their host relies models used for laboratory research. The closer a model matches the host, the more conclusive results are. Complex cell systems based on primary epithelial or stem cells are the gold standard of *in vitro* research. However, they are expensive, time consuming, and laborious to establish. Therefore, cell lines remain the backbone of virus research.

Despite vastly increased research into human coronaviruses following the COVID-19 pandemic, researchers continue to rely on suboptimal cell line models, for example ells of non-human origin like the VeroE6 African Green Monkey cell line. Using known expression patterns of the entry factors of the COVID-19 causative agent severe acute respiratory syndrome coronavirus-2 (SARS-CoV-2) we identified the Caki-1 cell line as a target for SARS-CoV-2. This cell line could be shown to be infectable with a wide range of coronaviruses including common cold virus hCoV-229E, epidemic virus MERS-CoV, and SARS-CoV-2 as well as other important respiratory viruses influenza A virus and respiratory syncytial virus. We could show that SARS-CoV-2 proteins can be stained for and localized in Caki-1 cells and the cells are competent of forming a cellular immune response. Together, this makes Caki-1 cells a unique tool for cross-virus comparison in one cell line.

## Introduction

Coronaviruses are a family of enveloped, positive-sense RNA viruses within the order *Nidovirales.* They are split into four genera: alpha-, beta-, delta-, and gamma-coronaviruses. *Alpha-* and *Betacoronaviruses* mainly infect mammalian hosts while the *Delta-* and *Gammacoronaviruses* primarily infect avian hosts [1]. Four members of the human coronaviruses (hCoV) cause mild, cold-like, upper respiratory infections: *Alphacoronaviruses* -229E and -NL63 and *Betacoronaviruses* -HKU1 and -OC43. On three occasions in the last 20 years, zoonotic transmission events of *Betacoronaviruses* lead to epidemic and pandemic disease outbreaks with higher severity pathologies: Severe acute respiratory syndrome coronavirus (SARS-CoV) in 2002-2003 [2]; Middle Eastern respiratory syndrome coronavirus (MERS-CoV) 2012-ongoing [3]; and most recently SARS-CoV-2, the causative agent of the coronavirus disease 2019 (COVID-19) pandemic 2019-ongoing [4]. The potential for future zoonotic events increases as human-animal reservoir interaction increases, for example through population growth, displacement due to climate change, deforestation, and finding alternative food sources.

Understanding coronavirus infections and host-pathogen interaction is paramount to developing novel antiviral strategies, and to prepare for future epidemics. This makes it imperative to find robust cell line models that are widely permissive to different coronaviruses allowing for direct comparison increasing result validity.

Important criteria for selecting model cell lines include species origin, tissue relevance, method of immortalization (i.e. presence of overexpressed viral oncogenes), genetic characterization and karyotype, versatility, and ease of culturing. A number of cell lines were identified as being naturally permissive to SARS-CoV-2, most notably Vero E6, Caco-2, and Calu-3 cells [5]. Overexpression of the main viral entry receptor, angiotensin-converting enzyme 2 (ACE2), sometimes aided by the overexpression of the proteolytic processing factor transmembrane protease, serine 2 (TMPRSS2) was shown to render non-susceptible cell lines permissive to SARS-CoV-2 infection. Although the overexpression cell lines then are permissive to infection, there are limitations and drawbacks to their use in research. Most notably, Vero E6 cells, which have been shown to be susceptible to many human viruses including influenza A virus (IAV), SARS-CoV, and MERS-CoV, are of African Green Monkey (*Chlorocebus aethiops*) origin, making them a less clinically relevant model. More importantly, they are deficient in type I interferon production due to a large deletion in their genome [6]. Tissue culture adaptation mutations and attenuation in the viral genome, such as the loss of the S1/S2 furin-cleavage site in SARS-CoV-2 spike (S), have been observed when the virus was passaged on Vero E6 cells [7]. Although the other notable cell lines commonly used to study SARS-CoV-2 are of human origin, viral titers obtained from Calu-3 and Caco-2 are lower than in Vero E6 cells [8]. Calu-3 and Caco-2 cells are heterogeneous populations of not fully differentiated cells, leading to varied consistency based on culturing methods. Some people also report difficulties culturing Calu-3 cells. Other, less explored cell lines were shown to be susceptible to high-titer SARS-CoV-2 infection, such as the breast cancer cell line CAL-51 [5],

The need for consistent and comparable cell lines has been highlighted by results from several genome-scale CRISPR knock-out studies investigating host factors during SARS-CoV-2 infection. The screens used different cell lines including Vero E6 [9], A549-ACE2 [10, 11], Huh7.5 [12], or a clone of Huh7.5 modified to overexpress both ACE2 and TMPRSS2 (Huh7.5.1-ACE2-IRES-TMPRSS2) [13]. Of these studies, ACE2 and cathepsin L (CTSL) were the only gene hits to overlap across all screens. The strongest overlap was observed between screens utilizing the same cell lines [10, 11] or clonal variants of a cell line [12, 13]. Although this suggests that there is reproducibility when using the same cell line, some cell lines modified for overexpression of cellular receptors have limited use for cross-comparison studies between viruses. For instance, while A549-ACE2 cells would be sufficient for comparing SARS-CoV and -2, both of which utilize ACE2, they could not be used to compare to other coronaviruses, such as hCoV-229E utilizing alanyl aminopeptidase (ANPEP) or MERS-CoV utilizing dipeptidyl peptidase 4 (DPP4) during entry. Additional overexpression or selection of clonal populations could change properties when comparing to other viruses. Huh7.5 cells and their derivatives are more broadly useable as they are naturally infectable by both hCoV-229E, and MERS-CoV due to their expressing a wide range of cell surface receptors. However, they are not very robust and exhibit extreme cytopathic effect upon early infection, limiting the types of studies to be conducted in these cells.

We aimed to find an adherent cell line that was both easy to transfect and naturally permissible to SARS-CoV-2. Both the human protein ATLAS and gene expression data deposited on NCBI identify the kidney as a tissue with high ACE2 expression. Pathology has furthermore identified SARS-CoV-2 infection within the kidneys of infected patients. Therefore, a human cell line isolated from a male clear cell carcinoma patient, Caki-1 (ATCC HTB-46), which has been previously used in cancer and toxicology studies was identified as a candidate target cell line. It has been well characterized through a number of loss-of-function screens and omics characterizations, as well as having data on drug sensitivities through the use of compound viability screens. A complete list of datasets can be found through the depmap portal (https://depmap.org/portal/cell_line/ACH-000433). Although Caki-1 cells have previously been reported to be permissive to two members of the family *Poxviridae*, myxoma virus [14] and Tanapoxvirus [15], we were unable to find reports of their being tested for infection with any RNA viruses.

Here, we describe the successful infection of Caki-1 cells when challenged with coronaviruses SARS-CoV-2, MERS-CoV, and hCoV-229E. All viruses replicated to similar or improved levels compared to commonly used cell lines. Using a SARS-CoV-2 anti-N antibody, we could show comparable replication kinetics between Caki-1 and Vero E6 cells. A panel of 26 antibodies raised in sheep against the known structural and non-structural proteins encoded by SARS-CoV-2 were used to study their cellular location during the SARS-CoV-2 replication in Caki-1 cells. We found good expression of virus receptors, interferon competence and susceptibility to gene knockdown in Caki-1 cells. Furthermore, two other clinically important respiratory RNA viruses, influenza A virus (IAV, order *Articulavirales*), and respiratory syncytial virus (RSV, order *Mononegavirales*) were also found to infect Caki-1 cells to comparable levels with the commonly used cell lines used for these viruses.

## Results

To compare the differences between infection in Caki-1 cells and the typical amplification cell line for each coronavirus, time courses of infection were performed for SARS-CoV-2, including variants of concern (VOCs), MERS-CoV, and hCoV-229E.

### Caki-1 cells show matching or improved infectability with SARS-CoV-2 compared to Vero E6 cells

Cells were infected with SARS-CoV-2 D614G or VOCs alpha, delta, or omicron (BA.1) to investigate how the growth profile differed between clinical isolates from early stages of the pandemic through to the current predominant variant, omicron (Results

A). Cells at confluence were inoculated with MOIs 0.1, 1, or 10 as determined by endpoint titration on Vero E6 cells, except for omicron, which was titrated on Caki-1 cells due to poor growth on Vero E6 cells. Following media replacement at 1.5 hours post inoculation (hpi), supernatant samples were collected and viral RNA quantities in the supernatant determined by direct lysis against a standard curve of known titer samples. The European “Original” strain EDB-2 showed a similar replication pattern in Caki-1 and Vero E6 cells at all MOIs. Both reached a plateau at 24 hpi at a relative TCID_50_/ml of ∼10^7^ and maintained that value until the 120 hpi time point. There was little difference in the log phase for EDB-2 between cell lines and MOIs, although the Vero E6 cells show greater variation across the time points compared to Caki-1 cells in which variation decreases as the MOI increases (Results

A). For the alpha variant, first detected in November 2020, we used strain EDB-α-1 for infection. The replication curve in Vero E6 cells flattens into a more linear phenotype peaking at 96 hpi with a relative TCID_50_/ml in the low 10^6^ region for all MOIs. The MOI of 0.1 infection lags slightly behind. Contrastingly, viral replication in Caki-1 cells still shows exponential growth, with stratification between the MOIs through the log phase before all plateauing at 48 hpi with a relative TCID_50_/ml maintained at ∼10^7^ (Results

A). For the delta variant, which emerged in late 2020, we used strain EDB-δ-1. The delta variant did not replicate as well as either previous variant overall and shows a more linear growth phenotype, similar to the EDB-α-1 variant in Vero E6, in both Vero E6 and Caki-1 cells. In both cell lines replication plateaus at 72 hpi up to 2-3×10^6^ TCID_50_/ml by 96 hpi. For EDB-δ-1, the increase in viral titer across the entire time course, in both cell lines, was only two logs. In contrast, in Caki-1 cells for EDB-2 and EDB-α1 the fold-change was five log_10_ (Results

A). The last VOC, omicron, was detected in December 2021 with new subvariants emerging since. Here, using EDB-ο-BA.1-10, we saw very poor infection in Vero E6 cells, barely increasing by 10-fold across 120 hpi, and dropping thereafter, likely due to virus degradation. In Caki-1 cells, we see a growth phenotype comparable to the original EDB-2 strain with a log phase peaking at 24 hpi and stratification between the different MOIs. Once EDB-ο-BA.1-10 reaches a plateau in Caki-1 cells at a TCID_50_/ml of ∼10^6^, it remains steady with very little variation for the remainder of the time course. Viral replication of EDB-ο-BA.1-10 in Caki-1 is reaching higher levels compared to Vero E6 cells (Results

A). As the virus evolved, it remained capable of replicating in the Caki-1 cells whilst progressively weaker infection is established in Vero E6 cells. The one exception is EDB-δ-1, which does not replicate well in either cell line.

### Caki-1 cells show improved infectability with MERS-CoV and hCoV-229E compared to Huh-7

Since Caki-1 cells showed good permissibility to infection with SARS-CoV-2, other coronaviruses were tested to determine if Caki-1 cells could be used as a pan-coronavirus model. To this end, the highly pathogenic beta coronavirus, MERS-CoV, and the less pathogenic common cold alpha coronavirus, hCoV-229E, were tested for replication in Caki-1 cells (Results

B+C). The traditional cell line for both these viruses is Huh-7.

For MERS-CoV infection, both Caki-1 and Huh-7 cells were inoculated using titres determined on Huh-7 cells. After 2 h, inoculum was replaced and samples taken at indicated time points. Infectious virus produced was assessed by endpoint titration on Huh-7 cells. In initially infected Huh-7 cells, the viral titer for MERS-CoV rose rapidly to 24 hpi and peaks here for an MOI of 1, but is slightly delayed for the lower MOIs, peaking at 48 hpi for 0.1 and 0.01 (Figure 1B). At all MOIs, a peak of ∼1.2×10^7^ TCID_50_/ml titer is achieved. After peaking, a rapid decline is observed for every MOI, crashing down to 10^4^ TCID_50_/ml by 96 hpi. Whilst the exponential phase in Caki-1 cells shows initial similarities to Huh-7 the viral titers plateau at 48 hpi for all MOIs. By that time, the cells have managed to support the virus to reach a TCID_50_/ml of ∼2.1×10^9^, which is two logs higher than Huh-7 cells in any condition. Whilst MOIs of 0.1 and 1 show a decreasing titer to ∼10^5^ TCID_50_/ml, MOI 0.01 infectious titers remain high up to 96 hpi. Caki-1 cells show a strong potential with MERS-CoV to be used as a new model system for growing the virus, higher titers are achieved with lower infection concentrations and the titers do not decrease with cell apoptosis as Caki-1 cells can robustly support the infection to its completion.

**Figure 1:**
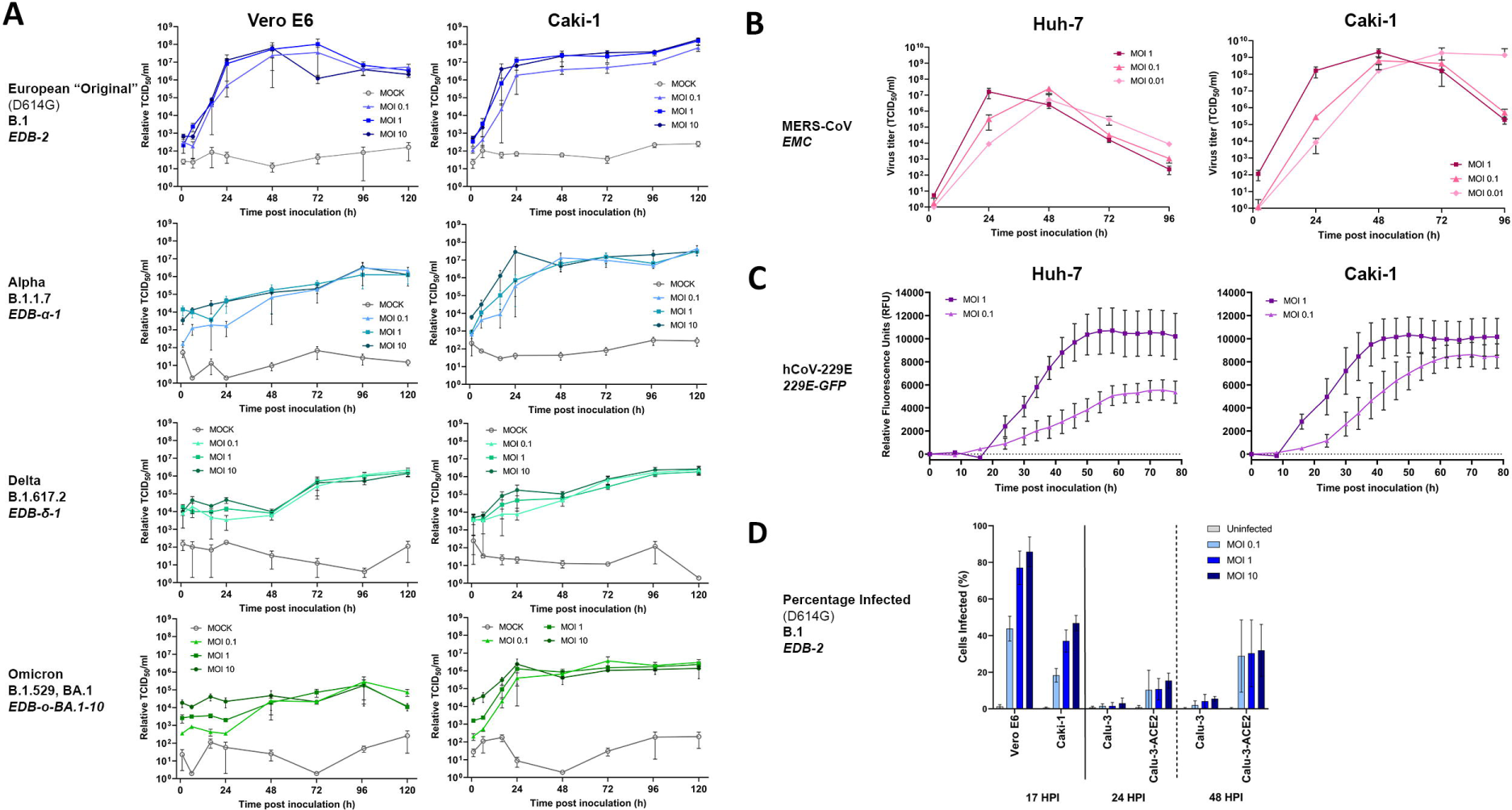
Time course assays in coronaviruses comparing the most commonly used cell line with Caki-1. For all graphs: circle points = MOI 10, square points = MOI 1, triangle points = MOI 0.1, diamond points = MOI 0.01, and the darker the color the higher the MOI, error bars indicate +/− SEM. [A] Time course infection of SARS-CoV-2 variants on Vero E6 and Caki-1 cells, biological n=3×3. Measured through RT-qPCR for the SARS-CoV-2 N gene in the supernatant, samples taken at 0, 6, 16, 24, 48, 72, 96 and 120 hpi at MOIs of 10, 1 and 0.1. [B] Time course infection of MERS-CoV on Huh-7 and Caki-1 cells, biological n=3×3, measured through TCID50 on Huh-7 cells. Samples taken at 24h intervals until 96hpi. [C] GFP time course of infection for hCoV-229E on Huh-7 and Caki-1 cells, measured from 0 hpi to 78 hpi in 6 h intervals, with fluorescence as a proxy for viral replication, biological n=3×3. [D] Fluorescence-activated cell sorting to determine the percentage of infected SARS-CoV-2 cells in Vero E6, Caki-1, Calu-3 and Calu-3-ACE2 cells. Vero E6 and Caki-1 cells were fixed at 17 hpi, stained for intracellular SARS-CoV-2 N protein and sorted for fluorescent cells, biological n=3×3. The same was conducted for Calu-3 and Calu-3-ACE2 cells at 24 hpi and 48 hpi, biological n=3×3.

To test whether Caki-1 cells are permissive to hCoV-229E, a GFP-reporter virus was used to measure infection throughout the course of infection using relative fluorescence of GFP as a proxy for viral replication rather than TCID_50_/ml (Results

C). Caki-1 and Huh-7 cells were inoculated at indicated virus concentrations as determined on Huh-7 cells. After 1.5 h, inoculum was replaced and plates returned to an incubating plate reader for analysis. The level of fluorescence achieved by both cell lines when infected is comparable to each other, but in Caki-1 cells, we see a quicker and more exaggerated log phase, reaching plateau at roughly 40 hpi, about 10 h before Huh-7 cells. There is also a difference in the lower MOI (0.1), which only achieves a plateau fluorescence value around 5,000 RFU in the Huh-7 cells, a >2-fold reduction compared to Huh-7 cells infected at an MOI of 1, while Caki-1 cells plateau around 8,500 RFU, only 2,000 RFU lower than the MOI of 1 and within the margin of error. The Caki-1 cells also show an observable difference of GFP distribution and brightness when compared to Huh-7 cells. Caki-1 cells produce a more uniform phenotype, with less background fluorescence, the GFP is expressed to a greater extent in infected cells, and it takes longer for the cells to round up and lift off the plate. Huh-7 cells generally express the GFP very weakly and in a more punctate fashion, lifting off the plate very early on in infection, making fluorescence readings difficult and inaccurate (data not shown).

### Only around 50% of Caki-1 cells get infected by SARS-CoV-2 showing potential of selection for even more susceptible subpopulations

To gain greater insight into the infection in Vero E6 cells compared to the more heterogeneous Caki-1 cells, we inoculated the cells at different MOIs (determined on Vero E6 cells) and assessed infection following a single-round infection at 17 hpi through staining for intracellular N protein and analysis by flow cytometry (Figure 1D). For all MOIs analyzed, the level of infected Caki-1 cells is roughly half of the infected Vero E6 cells. The Vero E6 population manages to reach up to ∼85.3% infected cells by 17 hpi at an MOI=10 and within the margin of error compared to MOI=1, whereas the Caki-1 reach an infection ∼46.8%. Another cell line used for SARS-CoV-2 research is Calu-3, an epithelial lung adenocarcinoma cell line with heterogeneous ACE2 expression. The percentage of infected cells remains low in the wild-type Calu-3 population, reaching 6% infection when inoculated at MOI=10 by 24 hpi, and only increasing by 0.8% after 48 hpi. Calu-3’s which have been enriched for high expression of ACE2 through staining, fluorescence activated cell sorting, and expansion, showed a marked increase in percentage of infection cells, reaching 21% at an MOI=0.1 at 24 hpi and roughly 50% at all MOIs at 48 hpi (Figure 1D). Whilst Calu-3 cells take longer to establish an infection, if enriched with ACE2 they are capable of reaching infection levels similar to Caki-1 cells although this is most likely through a multi-round infection unlike the single-round infection in Caki-1 cells at 17 hpi.

Caki-1 cells are permissible to all tested coronaviruses and in EDB-ο-BA.1-1, MERS-CoV and hCoV-229E, show a stronger infection than in Vero E6 and Huh7 cells, respectively. As SARS-CoV-2 VOCs emerged, Caki-1 cells have become a better infection model than the Vero E6 cells. They also provide a good replication model for other coronaviruses and can outperform the standard Huh-7 cells.

### Western blot analysis shows comparable rates of replication staining for the SARS-CoV-2 N protein

To gain a greater understanding of SARS-CoV-2 infection, antibodies were raised in sheep against all major proteins encoded by the SARS-CoV-2 genome (Figure 2A) by the University of Dundee MRC-Protein Phosphorylation Unit [16]. To assess the rate of replication at the protein level, viral replication was observed through western blot analysis against N protein of lysed cells. Briefly, Vero E6 and Caki-1 cells at confluence were inoculated at MOI=1 (as determined on Vero E6 cells) for 1.5h before media replacement. At indicated time points, cells were lysed and subjected to SDS-PAGE and western blot analysis. Quantification of protein levels normalized to beta-actin show comparable replication between Vero E6 cells and Caki-1 cells for EDB-2. As observed in the virus production levels, the replication of EDB-δ-1 was significantly slower with a first plateau reached at 16h, eventually increasing further to 48hpi but significantly lower in comparison to EDB-2 (Figure 2B). A background band could be observed even in uninfected cells at around 40kDa overlapping with the signal for beta-actin. This band was found to disappear when a rabbit anti-beta-actin antibody was used in later experiments (data not shown) indicating cross-reaction of the secondary anti-sheep or anti-mouse antibodies.

**Figure 2:**
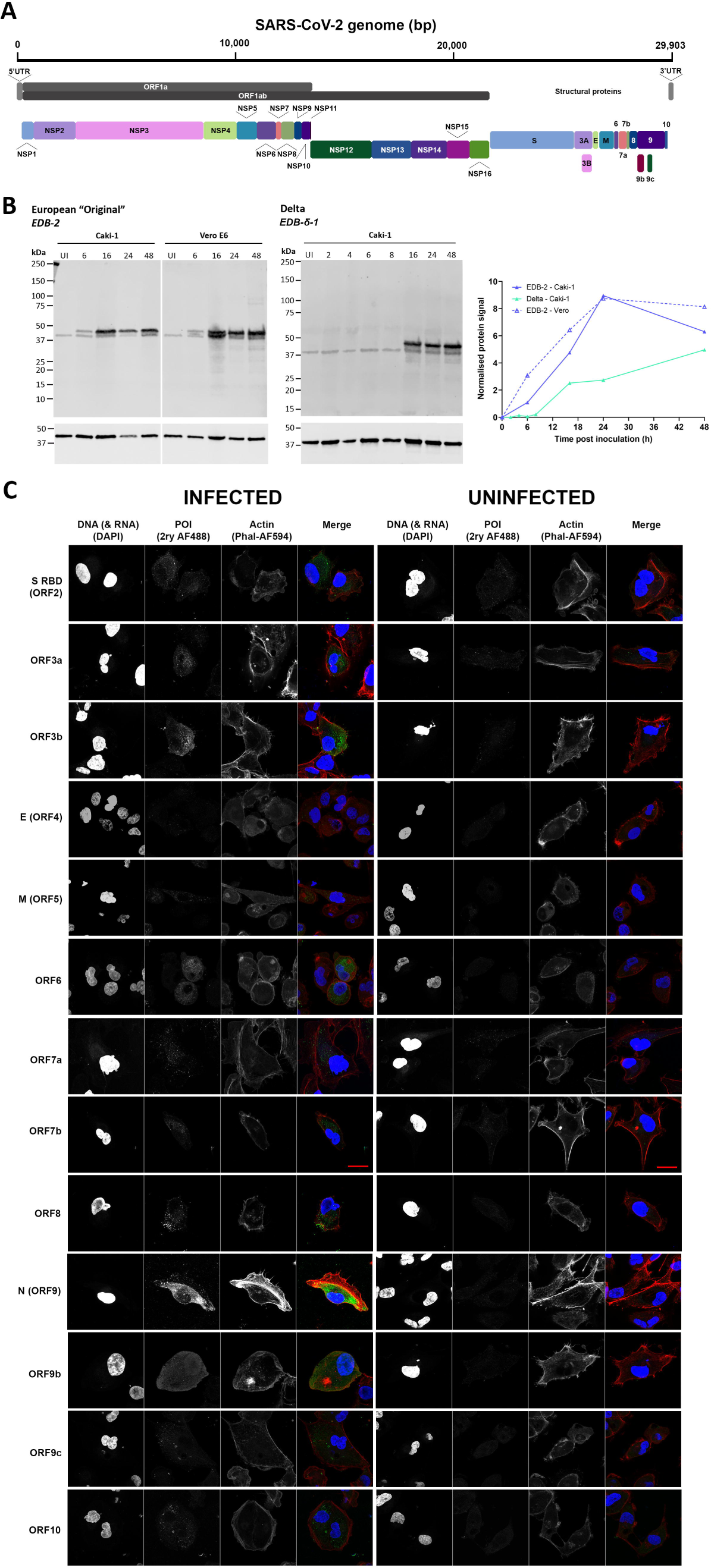
Confocal analysis of Caki-1 cells showing localization and size of SARS-CoV-2 orf proteins. [A] SARS-CoV-2 genome illustrating the proteins translated by the viral genome and their relative sizes. [B] Immunocytochemsitry staining showing infected and uninfected Caki-1 cells stained for SARS-CoV-2 open reading frame proteins. The protein of interest (POI) is indicated in green, the DAPI indicating DNA and RNA in blue, and actin in red. Scale bar represents 20 µM.

### Localization of viral proteins in Caki-1 cells during replication may pinpoint towards functionality

To fully characterize the SARS-CoV-2 replication in Caki-1 cells the full panel of 26 antibodies targeting both structural and non-structural proteins of the virus (as described above) were used to localize the proteins at peak infection. Caki-1 cells infected at a MOI=1 with SARS-CoV-2-EDB-2 were fixed at 16 hpi and stained with antibodies targeting most non-structural proteins and the known open reading frames in the “structural” section of the genome. A compromise was found using the antibodies, which had varying stock concentrations, at 1 in 200. Concentrations are therefore not comparable and for specific use of an antibody, we recommend further optimization. Further details may be found in the materials and methods section and table 1. Confocal images were acquired as z-stacks on a Zeiss LSM 880 (Airyscan) without use of the airyscan function. Imaging conditions were set for each uninfected, stained background cell with the matching primary antibody. Images depict a maximum projection of the z-stack.

**Table 1:**
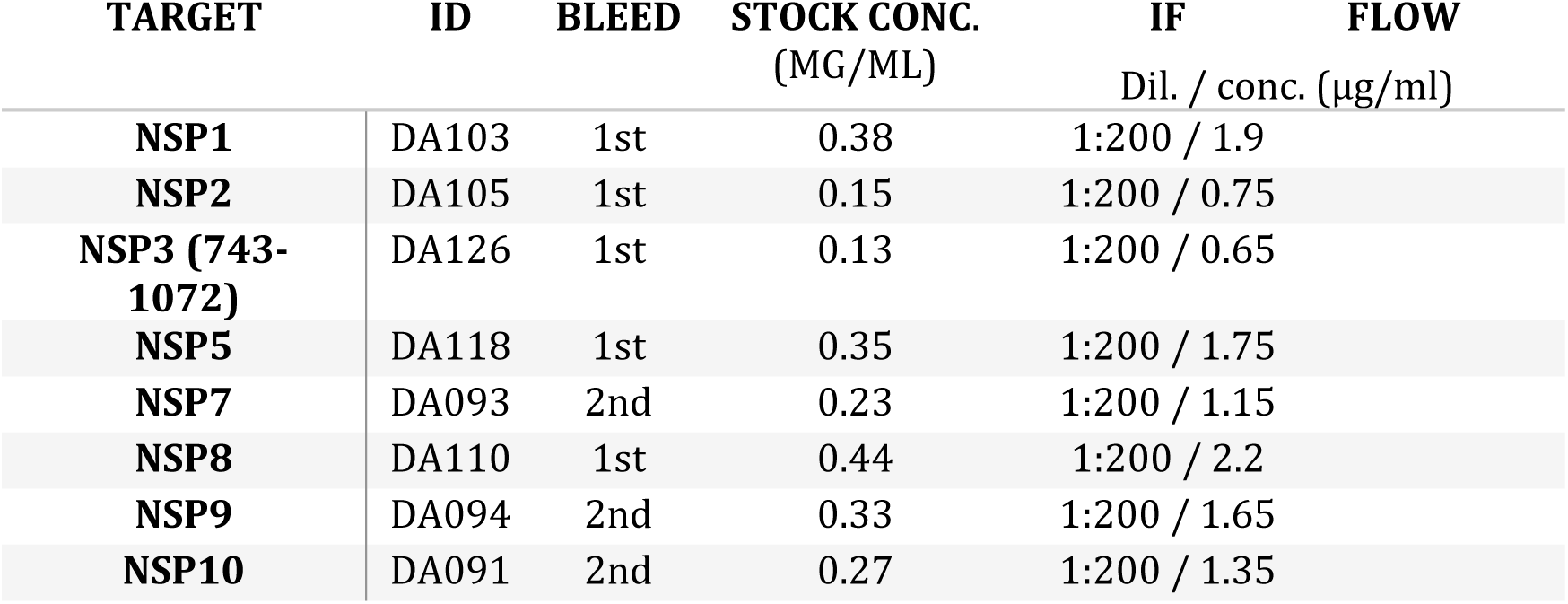
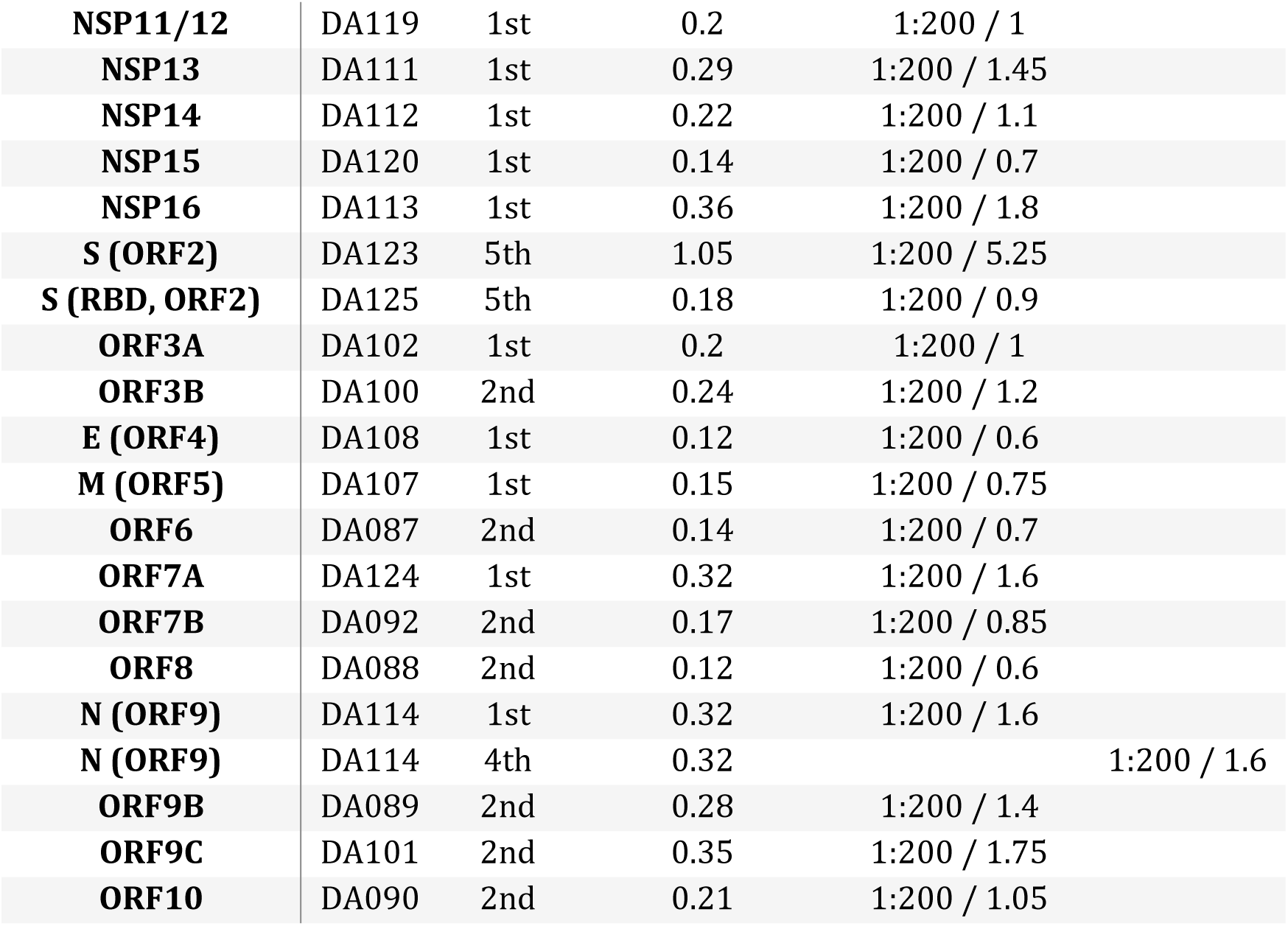
Antibodies targeting SARS-CoV-2 generated by the MRC PPU and respective concentrations used.

Of the open reading frame (ORF)2-10 encoded “structural” proteins, ORF3a and b, ORF6 ORF8 and N (ORF9) show the strongest signals above background. Almost all SARS-CoV-2 ORF antibodies show specific staining above background with the exception of E (ORF4) (Figure 2B). The spike receptor binding domain (S RBD), ORF2, shows vesicle-like punctate throughout the cytoplasm with a slight enrichment in the enlarged perinuclear space in some cells. There is little staining for S at the plasma membrane, which is in agreement with previous findings showing that only roughly 20% of coronavirus S protein may be found at the plasma membrane in the presence of M [17, 18]. In contrast to S, the M protein (ORF4) was found to be more perinuclear in localization and more punctate (Figure 2B). The viroporin ORF3a, shows similar though stronger punctate staining with concentration around the perinuclear space. This agrees with current SARS-CoV literature and reports of it being involved in autophagy, localizing to the lysosomes [19, 20]. Although there is some background staining in uninfected Caki-1 cells for ORF3a, the staining in the infected control is significantly stronger and background signal is non-specifically localized indicating non-specific background binding that may be optimized using different staining conditions. The small ORF3b protein, which acts as a potent interferon antagonist, localizes in the distal cytoplasm with some vesicular patterning. ORF3b also shows staining at the plasma membrane particularly along actin filaments. This agrees with reports for SARS-CoV of it being a multi-localized protein due to its shuttling behavior [21]. Co-localization studies with known interaction partners, such as IRF3, may shed further light into the role of ORF3b [22]. The interferon antagonist ORF6, binding and sequestering STAT1, is a known virulence factor of SARS-CoV-2, previously found to localize in the ER and at lysosomal membranes upon overexpression [23, 24]. A similar localization pattern is observed upon infection in Caki-1 cells with perinuclear concentration and distributed puncta throughout the cell. Additional staining could further substantiate localization to specific organelles (Figure 2B). SARS-CoV-2 ORF7a and ORF7b are type-I transmembrane proteins also involved in antagonism of the interferon response. Both are expressed and retained intracellularly. Background staining is observed for both these antibodies in the uninfected Caki-1 cells, but the fluorescence is stronger in the infected cells. ORF7a shows vesicular, diffuse localization, whilst ORF7b is concentrated in the perinuclear space with some plasma membrane staining. Little is known about the function of the two proteins, but for example ORF7b was shown to be incorporated into SARS-CoV particles [25]. The protein encoded by the SARS-CoV-2 ORF8 gene is unique to SARS-CoV-2 containing a predicted Ig-like fold, which can interact with a variety of host proteins involved in ER-associated degradation. Caki-1 cells show a strong staining with no background in the uninfected cells, and we see a vesicular staining with a concentration towards the plasma membrane in agreement with the existing literature showing secretion of ORF8 [26]. ORF9, the nucleocapsid protein (N), shows localization to the replication-transcription complex in the perinuclear space, where it aids packaging of the viral genome into a ribonucleoprotein particle. Staining for N is abundant across the cell, and even shows a small amount of nuclear staining (the seemingly stronger nuclear staining visible in the image is a consequence of maximum projection). N is also secreted into the extracellular space in larger amounts than ORF8, indicated by some of the vesicular and distal staining [27, 28]. An alternative ORF located within the N gene is ORF9b, which plays a role in suppressing innate immunity. ORF9b shows a very diffuse staining across the cell with no background in the uninfected cells. Reports have linked this protein with the mitochondrial membrane [29, 30], and upon close inspection, long punctate networks can be seen within some of the staining, although this is not obvious indicating mitochondrial interaction not to be a primary function of the protein. Most importantly, ORF9b shows clear nuclear localization. An second alternative ORF located within N encodes for ORF9c, a membrane-associated protein and has been associated with antiviral response suppression and suggested to be unstable [31]. Our results show an interesting concentration of ORF9c protein at the end of what look to be actin protrusions. Accumulations near the nucleus could however indicate an unfolded protein response (Figure 2B). ORF10 is possibly one of the most obscure proteins of SARS-CoV-2, with little known about its function. We found some perinuclear aggregations of the protein and otherwise relatively evenly distributed small puncta throughout the cell (Figure 2B).

The SARS-CoV-2 non-structural proteins (NSPs) are generated through proteolytic cleavage of the ORF1a and ORF1ab polyproteins. Of these, NSPs 3, 8, 10 and 16 gave the clearest staining. However, some NSPs showed very little specific staining, primarily NSP9 for which no staining could be seen in infected Caki-1 cells (Figure 3). The N-terminal cleavage products NSP1 and NSP2 are conserved across the coronavirus family and are both involved in halting host translation [32]. Whilst NSP1 shows very sparse staining in very distinct puncta throughout the cytosol, staining for NSP2 is concentrated around the perinuclear space. NSP3 is responsible for cleavage of NSP1, NSP2, and NSP3 from the N-terminal region of the polyproteins [33], and shows widespread, punctate staining. Some cells, possibly at an earlier stage in their infection cycle, show a more perinuclear focused staining, likely co-localizing with the replication transcription complex (RTC). The main proteinase NSP5 shows a very strong perinuclear, likely RTC, phenotype. NSP8 and NSP7 form a heterodimer. It is then unsurprising that staining for the proteins show similar localization. Whilst NSP7 is a relatively weak signal, NSP8 exhibits strong perinuclear staining showing big spots of protein staining, defined by its role in the RTC [34]. NSP8 also shows some nuclear staining, however, since there is some background nuclear staining it is unlikely this is a real signal (Figure 3). The growth-factor-like protein NSP10, which serves as a stimulatory factor for NSP14-ExoN [35], shows broadly distributed punctate staining with some localization to the plasma membrane and although the staining for NSP14 is not as strong, it demonstrates a similar pattern but with no membrane localization. NSP11_12, the latter being the RNA-dependent-RNA-polymerase, shows staining localizing to the RTC, as do the helicase NSP13, the endoribonuclease NSP15. There may be some staining for NSP13 in the nucleus. NSP16, a methyltransferase, shows nuclear staining and punctate staining throughout the cytosol indicating vesicular localization, and potentially some secretion [36] (Figure 3).

**Figure 3:**
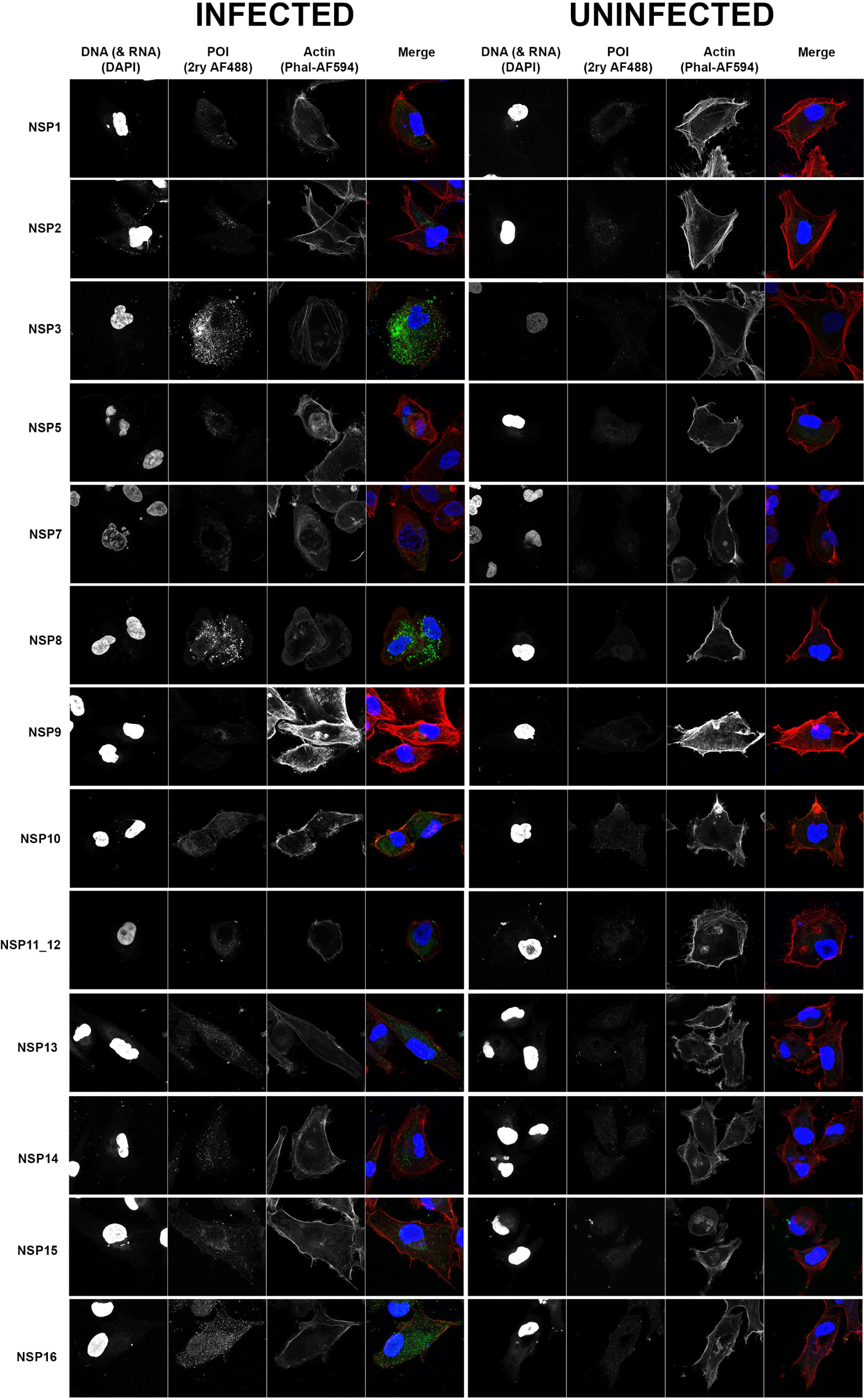
Confocal analysis of Caki-1 cells showing localisation and size of SARS-CoV-2 NSP proteins. Immunocytochemsitry staining showing infected and uninfected Caki-1 cells stained for SARS-CoV-2 non-structural proteins. The POI is indicated in green, the DAPI indicating DNA and RNA in blue, and actin in red. Scale bar represents 20 µM.

### Caki-1 cells show expression of receptors linked to SARS-CoV-2 and other human coronaviruses

To further understand the suitability of Caki-1 cells for infection assays and to account for the high permissibility to many coronaviruses, receptor expression profiling, interferon competence and siRNA knockdowns were conducted. The receptors/entry proteins for SARS-CoV-2, MERS-CoV and hCoV-229E, were checked for mRNA quantity by RT-qPCR, and normalized against a housekeeping gene (RRN18S) (Figure 1: **Time course assays in coronaviruses comparing the most commonly used cell line with Caki-1.** For all graphs: circle points = MOI 10, square points = MOI 1, triangle points = MOI 0.1, diamond points = MOI 0.01, and the darker the color the higher the MOI, error bars indicate +/− SEM. [A] Time course infection of SARS-CoV-2 variants on Vero E6 and Caki-1 cells, biological n=3×3. Measured through RT-qPCR for the SARS-CoV-2 N gene in the supernatant, samples taken at 0, 6, 16, 24, 48, 72, 96 and 120 hpi at MOIs of 10, 1 and 0.1. [B] Time course infection of MERS-CoV on Huh-7 and Caki-1 cells, biological n=3×3, measured through TCID50 on Huh-7 cells. Samples taken at 24h intervals until 96hpi. [C] GFP time course of infection for hCoV-229E on Huh-7 and Caki-1 cells, measured from 0 hpi to 78 hpi in 6 h intervals, with fluorescence as a proxy for viral replication, biological n=3×3. [D] Fluorescence-activated cell sorting to determine the percentage of infected SARS-CoV-2 cells in Vero E6, Caki-1, Calu-3 and Calu-3-ACE2 cells. Vero E6 and Caki-1 cells were fixed at 17 hpi, stained for intracellular SARS-CoV-2 N protein and sorted for fluorescent cells, biological n=3×3. The same was conducted for Calu-3 and Calu-3-ACE2 cells at 24 hpi and 48 hpi, biological n=3×3.

A). NRP1 (SARS-CoV-2 attachment), ANPEP (hCoV-229E receptor) and DPP4 (MERS-CoV receptor) all showed a similar level of relative expression in Caki-1 cells demonstrating that they are actively being expressed. Levels of around 1,000-fold lower expression than a housekeeping gene are in alignment with expression levels previously determined in lung (e.g. DPP4 [37]). ACE2 (SARS-CoV-2 receptor) and TMPRSS2 (one of the SARS-CoV-2 spike modifying proteases) had a relative lower amount of mRNA than the other factors, but this pattern correlates with expression of these proteins in nasal epithelial cells [38], perhaps providing an indication as to why they are so readily infectable with SARS-CoV-2.

### Caki-1 cells can be readily transfected with siRNA

To confirm Caki-1 cells are suitable cells in which to perform transfection experiments the effect of siRNA transfection targeting known entry factors of SARS-CoV-2 was examined by reading viral release 96 h post transfection (hpt) and 24 hpi (Figure 4B). A panel of genes implicated in SARS-CoV-2 entry (ACE2, TMPRSS2, NRP1, CTSB, CTSL), and a virus specific gene (SARS-CoV-2 N) were knocked down before infection with EDB-2 at 72 hpt. All genes showed a significant (****p<0.0001) reduction in replication compared to non-targeting siRNA, with SARS-CoV-2 N and NRP1 showing the strongest phenotypes with a relative replication compared to the non-targeting control of 8 and 9% respectively. Knockdown of CTSB had the least effect on viral replication, only causing a 50% reduction in released virus, followed by TMPRSS2 at 24.7% relative replication. The viability of the Caki-1 cells after knockdown was measured by fluorescence created through mitochondrial metabolism as a proxy for cell viability. There was no difference to cell viability after knockdown of any of these genes. Neither was there any effect on cell viability due to being treated with the transfection reagent (Mock). This shows that Caki-1 cells can be transfected successfully and that it results in the desired effect on viral replication, they can also tolerate the transfection reagent without a reduction in cell viability. Knockdowns were validated by RT-qPCR and showed at least 80% reduction in mRNA (data not shown).

**Figure 4:**
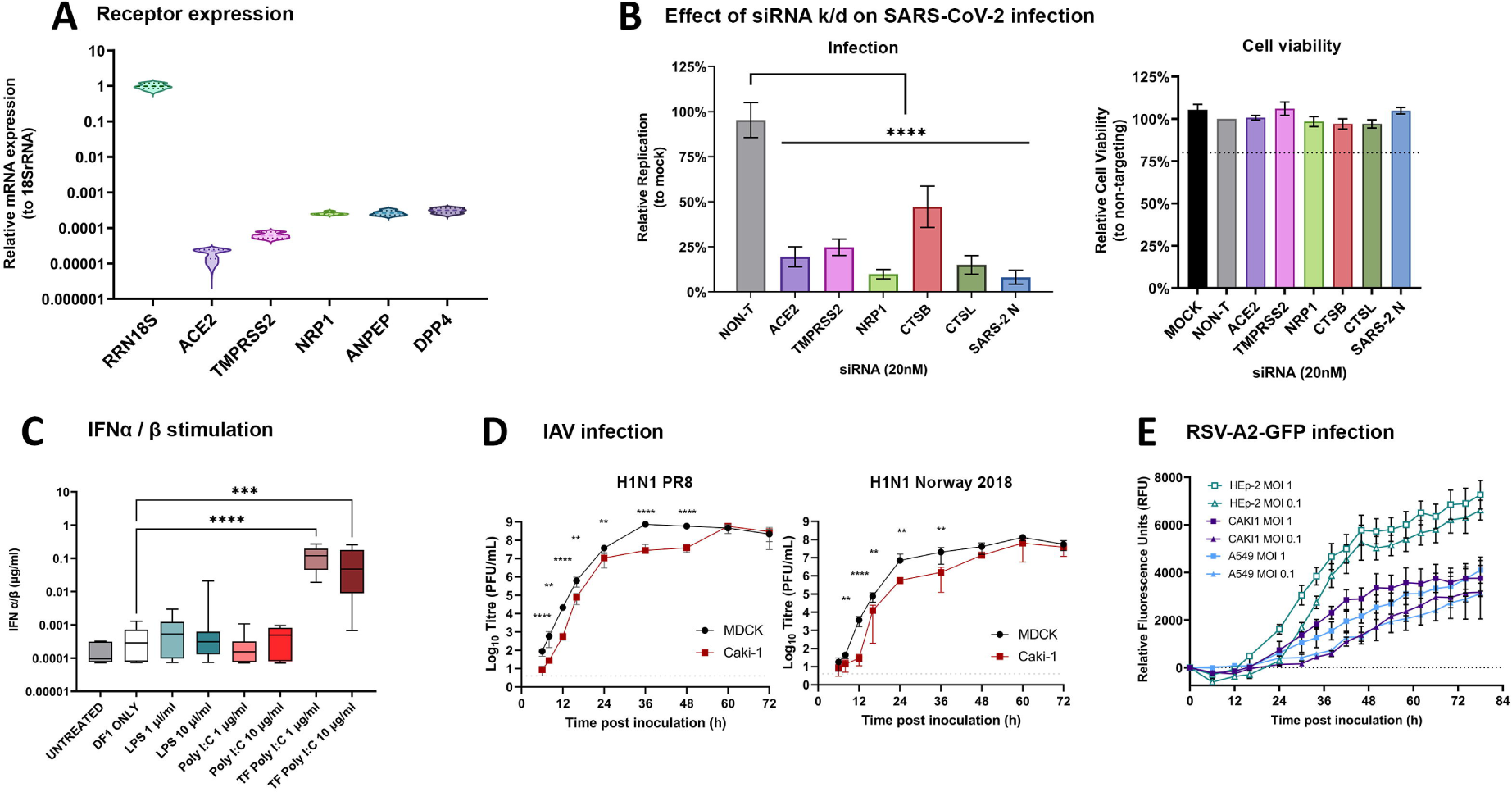
Characterizing Caki-1 cells in the context of respiratory virus infection with receptor and interferon competence profiling. [A] Violin plots showing mRNA expression of entry factors for the human coronaviruses ACE2, TMPRSS2, NRP1, ANPEP, and DPP4 in Caki-1 cells, relative to expression of the housekeeping gene RRN18S. Biological n=3×2. [B] (left) Relative replication of SARS-CoV-2 after knockdown of the entry factors in Caki-1 cells relative to the scrambled siRNA non-targeting at 24 hpi, all knockdowns are significant to a p-value < 0.0001 using a one-way ANOVA, biological n=12×3, error bars representing +/−SEM. (right) Cell viability after knockdown of SARS-CoV-2 entry factors for 72 h, dotted line indicates the cut-off for 80% relative cell viability, biological nx3, error bars represent +/−SEM. [C] Box and whiskers plot indicating the interferon α/β stimulation of Caki-1 cells after treatment with LPS and Poly I:C, TF indicates transfection into the cell. Transfection of Poly I:C at 1 µg/ml and 10 µg/ml were significant to 0.0001 and 0.001, respectively, using a one-way ANOVA. [D] Time course of IAV infection in Caki-1 cells (dark red filled squares) compared to MDCK (light red hollow squares) cells. Cells were infected at MOI 0.001 with either H1N1 PR8 or Norway 2018 strains as indicated, time points taken at 0, 12, 24 and 48 hpi and titered on MDCK cells, biological n=3, error bars representing +/−SEM. Significance was calculated using multiple Welch’s t-tests: *<0.05, **<0.01, ***<0.001, ****<0.0001. [E] RSV-A2-GFP replication in HEp-2 cells (turquoise hollow squares) compared to Caki-1 cells (dark blue filled squares) every 6 hpi up to 78 hpi monitored by fluorescence of GFP. Biological n=3×3, error bars represent +/−SEM.

### Caki-1 cells are interferon competent

Many cancer cells lines are deficient in their interferon response, especially Vero E6 cells, which lack the ability to produce interferon through a large deletion in their genome [6]. This can be detrimental for studying a fully comprehensive viral replication cycle, as close to natural infection as possible in a cancer cell line. It often also drives viral adaptation over serial passaging of field isolates. To assess the interferon response in Caki-1 cells, and therefore their suitability as a good infection model for SARS-CoV-2, we stimulated them with Poly I:C and LPS and measured their IFNα/β output by HEK-Blue IFNα/β assay (Figure 1: **Time course assays in coronaviruses comparing the most commonly used cell line with Caki-1.** For all graphs: circle points = MOI 10, square points = MOI 1, triangle points = MOI 0.1, diamond points = MOI 0.01, and the darker the color the higher the MOI, error bars indicate +/− SEM. [A] Time course infection of SARS-CoV-2 variants on Vero E6 and Caki-1 cells, biological n=3×3. Measured through RT-qPCR for the SARS-CoV-2 N gene in the supernatant, samples taken at 0, 6, 16, 24, 48, 72, 96 and 120 hpi at MOIs of 10, 1 and 0.1. [B] Time course infection of MERS-CoV on Huh-7 and Caki-1 cells, biological n=3×3, measured through TCID50 on Huh-7 cells. Samples taken at 24h intervals until 96hpi. [C] GFP time course of infection for hCoV-229E on Huh-7 and Caki-1 cells, measured from 0 hpi to 78 hpi in 6 h intervals, with fluorescence as a proxy for viral replication, biological n=3×3. [D] Fluorescence-activated cell sorting to determine the percentage of infected SARS-CoV-2 cells in Vero E6, Caki-1, Calu-3 and Calu-3-ACE2 cells. Vero E6 and Caki-1 cells were fixed at 17 hpi, stained for intracellular SARS-CoV-2 N protein and sorted for fluorescent cells, biological n=3×3. The same was conducted for Calu-3 and Calu-3-ACE2 cells at 24 hpi and 48 hpi, biological n=3×3.

C). Stimulation with a transfection reagent only (DF1) induced a small response, comparable to cells cultured in media containing Poly I:C. Caki-1 cells only showed a very small response to LPS stimulation, comparable to other stimulant addition. However, when Poly I:C was transfected to mimic a viral infection, IFN α/β production rose dramatically from 0.0004 µg/ml for DF1 only to 0.2 µg/ml for 1 µg/ml Poly I:C (****p<0.0001) and 0.08 µg/ml for 10 µg/ml Poly I:C (***p<0.001). None of the other conditions provided a significant difference from untreated. This demonstrates that Caki-1 cells can mount a competent interferon response, and so, in assays which investigate interferon pathways in viral replication, and any other host-pathogen interactions, they would provide a complete picture of viral replication.

### Caki-1 cells are broadly susceptible to infection with other respiratory viruses RSV and IAV

To determine whether Caki-1 cells are permissible to infection with other respiratory viruses not in the coronavirus family, they were infected with either the IAV H1N1 vaccine strain A/Puerto Rico/8/1934 (H1N1 PR8), a more recent H1N1 isolate A/Norway/3433/2018 (H1N1 Norway 2018), or RSV and compared to the standard cell line used to replicate these viruses. In a trypsin-dependent infection with both H1N1 strains, Caki-1 cells supported virus replication comparably to MDCK cells when infected with an MOI=0.001. No infection was observed in a trypsin-independent infection (data not shown). Slightly lower levels of infectious virus were found in Caki-1 cells up to 24 and 48 hpi for H1N1 Norway and PR8, respectively (Figure 1: **Time course assays in coronaviruses comparing the most commonly used cell line with Caki-1.** For all graphs: circle points = MOI 10, square points = MOI 1, triangle points = MOI 0.1, diamond points = MOI 0.01, and the darker the color the higher the MOI, error bars indicate +/− SEM. [A] Time course infection of SARS-CoV-2 variants on Vero E6 and Caki-1 cells, biological n=3×3. Measured through RT-qPCR for the SARS-CoV-2 N gene in the supernatant, samples taken at 0, 6, 16, 24, 48, 72, 96 and 120 hpi at MOIs of 10, 1 and 0.1. [B] Time course infection of MERS-CoV on Huh-7 and Caki-1 cells, biological n=3×3, measured through TCID50 on Huh-7 cells. Samples taken at 24h intervals until 96hpi. [C] GFP time course of infection for hCoV-229E on Huh-7 and Caki-1 cells, measured from 0 hpi to 78 hpi in 6 h intervals, with fluorescence as a proxy for viral replication, biological n=3×3. [D] Fluorescence-activated cell sorting to determine the percentage of infected SARS-CoV-2 cells in Vero E6, Caki-1, Calu-3 and Calu-3-ACE2 cells. Vero E6 and Caki-1 cells were fixed at 17 hpi, stained for intracellular SARS-CoV-2 N protein and sorted for fluorescent cells, biological n=3×3. The same was conducted for Calu-3 and Calu-3-ACE2 cells at 24 hpi and 48 hpi, biological n=3×3.

D). HEp-2 cells are typically used to expand RSV. Caki-1 and HEp-2 cells were inoculated with a fluorescence reporter virus RSV-A2-GFP. Replication was measured as a proxy by measuring the florescence reporter in an incubating plate reader. For the first time, the replication in Caki-1 cells does not match or outperform replication of RSV in HEp-2 cells (Figure 1: **Time course assays in coronaviruses comparing the most commonly used cell line with Caki-1.** For all graphs: circle points = MOI 10, square points = MOI 1, triangle points = MOI 0.1, diamond points = MOI 0.01, and the darker the color the higher the MOI, error bars indicate +/− SEM. [A] Time course infection of SARS-CoV-2 variants on Vero E6 and Caki-1 cells, biological n=3×3. Measured through RT-qPCR for the SARS-CoV-2 N gene in the supernatant, samples taken at 0, 6, 16, 24, 48, 72, 96 and 120 hpi at MOIs of 10, 1 and 0.1. [B] Time course infection of MERS-CoV on Huh-7 and Caki-1 cells, biological n=3×3, measured through TCID50 on Huh-7 cells. Samples taken at 24h intervals until 96hpi. [C] GFP time course of infection for hCoV-229E on Huh-7 and Caki-1 cells, measured from 0 hpi to 78 hpi in 6 h intervals, with fluorescence as a proxy for viral replication, biological n=3×3. [D] Fluorescence-activated cell sorting to determine the percentage of infected SARS-CoV-2 cells in Vero E6, Caki-1, Calu-3 and Calu-3-ACE2 cells. Vero E6 and Caki-1 cells were fixed at 17 hpi, stained for intracellular SARS-CoV-2 N protein and sorted for fluorescent cells, biological n=3×3. The same was conducted for Calu-3 and Calu-3-ACE2 cells at 24 hpi and 48 hpi, biological n=3×3.

E). However, the infection in Caki-1 cells outperforms RSV infection in A549 cells through faster and stronger (higher fluorescence) growth. The fluorescence from the HEp-2 cells (turquoise) reaches 7,000 RFU for MOIs of 1 and 0.1, with 0.1 remaining slightly under MOI=1 for the entire replication cycle (0 hpi – 78 hpi). Although the Caki-1 cells do show a logarithmic replication curve at both MOIs, it is shallow and plateaus by 42 hpi, about 10 h before the HEp-2 cells. The maximum RFU achieved by the hRSV-A2-GFP is ∼3,700, about half of the RFU achieved in HEp-2. The variation at the later timepoints has increased compared to HEp-2 cells. No syncytia formation was observed in the Caki-1 cells upon hRSV-A2-GFP infection, but CPE was present at 48 hpi.

## Discussion

Here we have shown that Caki-1 cells can be used to model infection for a range of respiratory viruses; SARS-CoV-2, CoV-229e, MERS-CoV, IAV, and RSV. With the exception of RSV, they are comparable to or outperform the current preferred cell line for viral production, whilst also providing a naturally permissible environment for the virus to replicate in. They express all the receptors needed for infection with SARS-CoV-2, MERS-CoV, and hCoV-229E, and are responsive to transfection with siRNA without significant cell death. They have a competent interferon response as demonstrated by stimulation with Poly I:C and are also permissive to other respiratory viruses. This makes them a highly useful model for direct comparison of respiratory viruses across one, unmodified (no overexpression) cell line.

The natural susceptibility of Caki-1 (human) cells to a range of respiratory viruses, although not a lung cell line, provides them with an advantage over many other cell lines used for viral research. Particularly over the much used Vero E6 cells (African Green Monkey), which, although permissive to many different viruses, do not provide a representative environment for human pathogens. The broad susceptibility of Caki-1 cells to respiratory virus infection allows for their use in comparative studies between viruses. Calu-3 cells, which have been widely used as a lung cell model for SARS-CoV-2, show a very low percentage infection and are difficult to grow. Selection for ACE2 expressing cells (Calu-3-ACE2) through FACS makes the cells more permissive to infection with SARS-CoV-2 and can also enhance growth characteristics through selection of cell subpopulations. However, this specializes the Calu-3-ACE2 cells to infection with SARS-CoV-2 and is likely impacting infectability with other viruses. Similarly though, if someone wanted a more highly permissive version of Caki-1cells to study their SARS-CoV-2 infection, selection of ACE2 cells may improve infection beyond 50%. We would however advise to keep this population as is due to its broad susceptibility to a range of respiratory viruses.

Whilst Caki-1 are still a cancer cell line, and thus contain the mutations and caveats associated with that fact, they have a well-characterized genome and expression patterns. Whilst they have a knockout mutation in the CDKN2 gene and this should be kept in mind, they also boast a fully receptive innate immune response, allowing this pathway to be interrogated (https://www.cbioportal.org/, Accessed 8/10/2022, [39, 40]). This is also demonstrated by their response to intracellular Poly I:C (a viral mimic), but not to LPS which is a bacterial stimulant of immune cells, showing their response is specific and relevant to viral infections. Conversely, the Vero E6 cells are defective in its production of interferon, but remains sensitive to the action of interferon through ISGs, often making it a preferred cell line for the production of viral stocks, with the caveat that attenuation is more likely [6].

With the emergence of new SARS-CoV-2 variants it has been found that in other cell lines, there is a change in tropism and attachment / processing factor preference leading to a loss of infection [41]. Caki-1 cells are able to maintain equal levels of infection across all the VOCs tested (alpha, delta, and omicron), without a change in phenotype, whereas Vero E6 cells lose their ability to effectively support viral replication as the variants evolve. This decrease in permissibility of the Vero E6 cells runs in parallel with the decrease in severe symptoms seen with the evolution of the variants, with omicron causing the mildest disease of all the variants, likely attributed to a change in cell tropism and / or temperature [42]. As the Caki-1 cells can support infection of the omicron variant, and it has been reported that omicron has moved away from TMPRSS2 mediated entry [43], it could be assumed that entry into Caki-1 cells is less or not TMPRSS2 dependent. This is reflected in the knockdown phenotypes (Figure 4B) with 25% relative infection still occurring upon knockdown of TMPRSS2. This makes the Caki-1 cell line very valuable for future SARS-CoV-2 research as they will be useful to study the host-virus interactions of variants evolved from omicron.

The Caki-1 cell line also offers ideal growth characteristics for viral study and propagation. Whilst being permissive to transfection Caki-1 cells have a doubling time of 36 hours (preventing overgrowth), the cultures can pack tightly to form dense monolayers capable of supporting widespread viral infection, and are robust enough to be able to maintain multi rounds of infection without significant CPE development. Conversely, Vero E6, Calu-3, and Huh7 cells all develop CPE after 24 hpi with SARS-CoV-2, restricting their use [7, 44]. The slower growth of Caki-1 cells compared to all other preferred cell lines, especially Vero E6 cells, also means that they are suitable for producing stocks of slower growing viruses such as the common cold coronaviruses. Here, we have also utilized Caki-1 cells to isolate clinical samples of SARS-CoV-2, specifically omicron, a methodology which usually has been achieved on Vero E6 cells for all major virus types. This negates the worry for mutation accumulation in the S1/S2 furin cleavage site that has been observed in SARS-CoV-2 within a single passage in Vero E6 cells [7]. The ability of Caki-1 cells to support robust infection with multiple respiratory viruses shows that this cell line can be useful to study both coronaviruses and common cold viruses in direct comparison. The growth properties of the cells furthermore improve high-throughput readouts and methods through slower growth and overcrowding, reducing background fluorescence and changes to cell properties in a dense cell layer.

The SARS-CoV-2 antibodies developed by MRC PPU Dundee form a useful toolset to quantify and visualize viral proteins throughout the replication cycle as we have demonstrated here. We have shown the localization of most viral proteins within Caki-1 cells, which agrees with the literature for other cell lines localization and relevant SARS-CoV homologues [45, 46], and tagged proteins in A549 cells [47]. Contrast to the studies previously performed, here we can show how natural expression of these proteins is characterized in cells where all other viral proteins and cellular interaction partners are regulated through the viral infection. The staining and the advantage of higher resolution techniques show the vast potential to study viral protein function through localization. Particularly interesting will be the function of nuclear localization of NSP16 and ORF9b in particular. The many functions associated with proteins like N, ORF3B, NSP2, and NSP8 are indicated through their broad distribution patterns within the cell. With these antibodies being raised in sheep, they are ideal for pairing with other antibodies for co-localization studies. Here, we have used a variety of antibodies purified from different bleeds of the production sheep as they were available to us. Indications are that later bleeds show less background and more specific staining. It is clear that further optimization for some of the staining protocols is warranted for some of the proteins of interest.

The multifaceted uses of Caki-1 cells in viral research are clear; they are easy to work with and manipulate in the lab, they can support a range of viruses across multiple rounds of infection, and in most cases outperform the commonly used cell line for each virus. They naturally highly express a wide range of receptors, and possess an interferon response that can both produce and respond, making them an ideal candidate for studying viral infections *in vitro*.

## Methods

### Cells & Viruses

Vero E6 (ATCC CRL-1586), Calu-3 (ATCC HTB-55), Huh-7 (JCRB, JCRB0403), HEp-2 (ATCC CCL-23), and MDCK (ATCC CCL-34) cells were maintained as monolayer cultures in Dulbecco’s Modified Eagle’s Medium (DMEM, Sigma), supplemented with 10% heat inactivated Fetal Bovine Serum (FBS, Gibco), 1X Ultraglutamine-I (Lonza), and 1X Non-essential Amino Acids (NEAA, Lonza) (complete DMEM). Calu-3 cells were supplemented with 1X Sodium Pyruvate (Sigma). Hek-Blue IFN-α/β (InvivoGen) cells were maintained as monolayer cultures in complete DMEM supplemented with 30 µg/ml of blasticidin (Gibco) and 100 µg/ml of Zeocin (Gibco). Caki-1 cells (ATCC HTB-46) were maintained as monolayer cultures in Roswell Park Memorial Institute medium (RPMI, Sigma), supplemented with 10% heat inactivated FBS (Gibco), 1X Ultraglutamine-I (Lonza), and 1X NEAA (Lonza). Cells were maintained at 37°C in 5% CO_2_.

Samples from confirmed COVID-19 patients were collected by trained healthcare professionals using combined nose-and-throat swabbing. The samples were stored in virus transport medium and sterile filtered through 0.1 μm filters prior to cultivation and isolation on Vero E6 cells. Variant B.1.1.529 (ο) was isolated on Caki-1 cells. Samples were anonymized by coding, compliant with Tissue Governance for the South East Scotland Scottish 279 Academic Health Sciences Collaboration Human Annotated BioResource (reference no. SR1452). Virus sequence was confirmed by Nanopore sequencing according to the ARCTIC network protocol (https://artic.network/ncov-2019), amplicon set V3, and validated against the patient isolate sequence. SARS-CoV-2 variants utilized in this study are EDB-2 (B1.5 at the time, now B.1), EDB-α-1 (B.1.1.7), EDB-δ-1 (B.1.617.2), and EDB-ο-BA.1-1 (B.1.1.529, BA.1). Infectivity was quantified by endpoint titration on Vero E6 cells, for all isolates apart from ο (omicron), which was titrated on Caki-1 cells. All infections in this study were performed using passage 2 from isolation of each variant.

hCoV-229E-GFP [48] was propagated on Huh7 cells at 34°C with 5% CO_2_ over 72 hours. Infectivity was quantified by endpoint titration on Huh7 cells.

hRSV-A2-eGFP [49] (obtained from Juergen Schwarze, University of Edinburgh) was amplified on HEp-2 cells over 72 hours. Infectivity was quantified by endpoint titration on HEp-2 cells.

MERS-CoV strain EMC [50] was propagated in VeroB4 cells. Infectivity was quantified by endpoint titration on Huh7 cells as previously described [51].

Influenza strain A/Puerto Rico/8/1934 H1N1 (PR8) was generated by reverse genetics as previously described [52]. A/Norway/3433/2018 H1N1 (Norway H1N1) was kindly supplied by the Worldwide Influenza Centre at the Francis Crick Institute, London, made from clinical samples they received from the WHO National Influenza Centre, Department of Virology, Folkehelseinstituttet, Oslo. A working stock was generated in MDCK cells.

### Viral Growth Curves

#### SARS-CoV-2

Vero E6 and Caki-1 cells were seeded to confluence in a 24-well plate one day prior to inoculation at differing MOIs for 1 h. At the indicated time point post infection, supernatant was lysed and quantified as previously described [53]. Briefly, the lysate was added to the reaction at 10% total reaction volume and analyzed by RT-qPCR, using SYBR Green GoTaq 1-Step RT-qPCR (Promega) with the SARS-CoV-2 CDC N3 primers (F – GGGAGCCTTGAATACACCAAAA, R –TGTAGCACGATTGCAGCATTG) at 350 nM each to determine viral copy number according to the manufacturer’s instructions (annealing at 60°C) on a Stratagene MX3000p qPCR system. To assess viral copy numbers, resulting Cts were analyzed against a standard curve of lysate with known TCID_50_/ml value as well as an RNA template.

#### hRSV and hCOV-229E

HEp-2 (RSV), Huh7 (229E), and Caki-1 (both) cells were seeded to confluence in a black, clear bottom 96-well plate one day prior to infection. After inoculation at differing MOIs for 1.5 h, inoculum was removed and replication measured as a function of GFP fluorescence from 6 hpi to 72 hpi in 2-hour intervals (BMG Clariostar; excitation: 470-15, dichroic: 492.5, emission: 515-15, bottom optic). Slopes of the linear phase of replication were calculated then normalized to uninfected cells (slope = 1) and the average slope calculated.

#### MERS-CoV

Huh7 and Caki-1 cells were seeded to confluence one day prior to infection. After inoculation at differing MOIs for 2 h inoculum was removed and cells were washed three times with PBS. At indicated time points post infection, supernatant was lysed and quantified as previously described. Virus titers were determined by endpoint titration on Huh7 cells inoculated with serial dilution and TCID_50_/ml was visualized using Crystal Violet 72 hpi and calculated by the Spearman–Kärber algorithm.

#### IAV

MDCK and Caki-1 cells were seeded to confluence on the day of infection. Once counted, cells were washed with PBS and infected at MOI 0.001. After a 1h adsorption, inoculum was replaced with serum-free medium supplemented with 1 µg/ml tosyl phenylalanyl chloromethyl ketone (TPCK)-treated trypsin (Sigma) and 0.14% BSA fraction V (w/v) (Sigma). Cells were incubated at 37°C and supernatant samples were harvested at the indicated time points.

Virus titers were determined by plaque assays in MDCK cells. Briefly, confluent monolayers of MDCK cells were washed with PBS, infected with ten-fold serial dilutions of virus samples performed in serum-free media and incubated at 37°C for 1h. Inoculum was replaced with DMEM including 0.14% BSA (Sigma), 1 µg/ml TPCK-treated trypsin (Thermo) and 1.2% Avicel (Merck). Following a 3-day incubation, overlay was removed and cells were fixed with 4% formaldehyde in PBS and subsequently stained with 0.1% Toluidine Blue (Sigma-Aldrich). Stained plates were washed under tap water and left to dry. Plaques were counted and viral titers were expressed as PFU/ml.

### Fluorescent Activated Cell Sorting (FACS) of Calu-3 cells

Calu-3 cells were enriched for high ACE2 expression (Calu-3-ACE2) by FACS using a BD Biosciences FACSAria III Cell Sorter. Cells were detached from a T75 using Gentle Cell Dissociation Reagent (Stemcell Technologies) and recovered in DMEM prior to staining. Recombinant Anti-ACE2 antibody (Abcam, ab272500) was used at 1:500 before staining with Goat anti-Rabbit IgG (H+L) AF647 (Invitrogen, A-21244) at 1:2000. The top 0.1% of cells expressing ACE2 were collected and cultured in conditioned media to recover and expand.

### Flow Cytometry

Vero E6, Caki-1, Calu-3, and Calu-3-ACE2 cells were seeded one day prior to inoculation with SARS-CoV-2-EDB-2 at differing MOIs of 10, 1 and 0.1 for 1 h. At the indicated timepoints, the cells were detached from the well using TrypLE Express Enzyme (Gibco) prior to fixation with 4% formaldehyde. The cells were permeabilized using 0.1% Triton-X-100. Using standard immunocytochemistry techniques, primary antibody targeting SARS-CoV-2 N protein (DA114 4^th^ bleed, MRC PPU Dundee, generated as described in [16]) was used at 1:200 (Table 1) before staining with Donkey anti-Sheep AF488 Secondary Antibody (Invitrogen, A-11015) at 1:2000. The cells were analyzed using a B530/30-A filter on a BD Biosciences LSRFortessa X-20 Cell Analyzer.

### Confocal Imaging

Caki-1 cells were seeded in clear-bottom Lumox plates (Sarstedt) one day prior to infection with EDB-2 at MOI = 1. At 16 hpi the cells were fixed with 4% formaldehyde and permeabilized using 0.1% Triton-X-100. Using standard immunocytochemistry techniques, primary antibodies targeting SARS-CoV-2 proteins were used at indicated dilutions / concentrations (Table 1, MRC PPU Dundee) before staining with Donkey anti-Sheep AF488 Secondary Antibody (Invitrogen) at 1:2000, DAPI (Invitrogen, A-11015) at 0.2 µg/ml, and Alexa Fluor™ 594 Phalloidin (Invitrogen) at 1:5,000. The cells were analyzed on a Zeiss LSM 880 Airyscan microscope acquiring z-stack images with a set 0.5 µm imaging interval in the z-direction. Imaging properties were set for an uninfected, stained control sample for each antibody. Images were processed using Fiji (ImageJ) to generate maximum projection images, merge, and montage.

### Receptor Quantification

RNA was extracted from Caki-1 cells using the RNeasy (Qiagen) kit according to the manufacturer’s instructions. RNA was quantified by Nanodrop and Qubit (ThermoFisher). RT-qPCR reactions using the GoTaq 1-Step RT-qPCR kit (Promega) were set-up to 10 µl final reaction volumes according to the manufacturer’s instructions containing 250 nM of each receptors primer pairs and 10 ng of RNA. The RT-qPCR was run on a Mx3000P (Agilent) using an annealing temperature of 60°C and analyzed against a standard curve of Mitochondrial 18S ribosomal RNA (RRN18S) before being normalized to the RRN18 value. Primers used were ACE2 (fwd: TTCCACTCTCATTTGAGCCTG, rev: GCCGGAGATAGGAGTGGA), TMPRSS2 (fwd: AGGGAAGACCTCAGAAGTGC, rev: CACAGATCATGGCTGGTGTG), NRP1 (fwd: TCCTCATCGGGCATTCTCTC, rev: TCTGAGACACTGCTCTGCAA, ANPEP (fwd: TTCAACATCACGCTTATCCACC, rev: AGTCGAACTCACTGACAATGAAG), DPP4 (fwd: TACAAAAGTGACATGCCTCAGTT, rev: TGTGTAGAGTATAGAGGGGCAGA), and RRN18S (fwd: AGAAACGGCTACCACATCCA, rev: CACCAGACTTGCCCTCCA).

### Receptor siRNA knockdown

siRNA (Dharmacon – Table 2) and Dharmafect 1 (Horizon) were combined in OptiMEM (Gibco) to achieve a final concentration of 20 nM and 0.1% respectively. Transfection complexes were incubated for 20 minutes at room temperature before the addition of 20,000 cells per 96-well. At 72 h post transfection, cells were infected with EDB-2 at MOI = 0.1. At 24 and 48 hpi, supernatant was harvested, lysed, and quantified as as previously described [53]. Relative quantities were normalized to the Non-targeting control (#1).

**Table 2:**
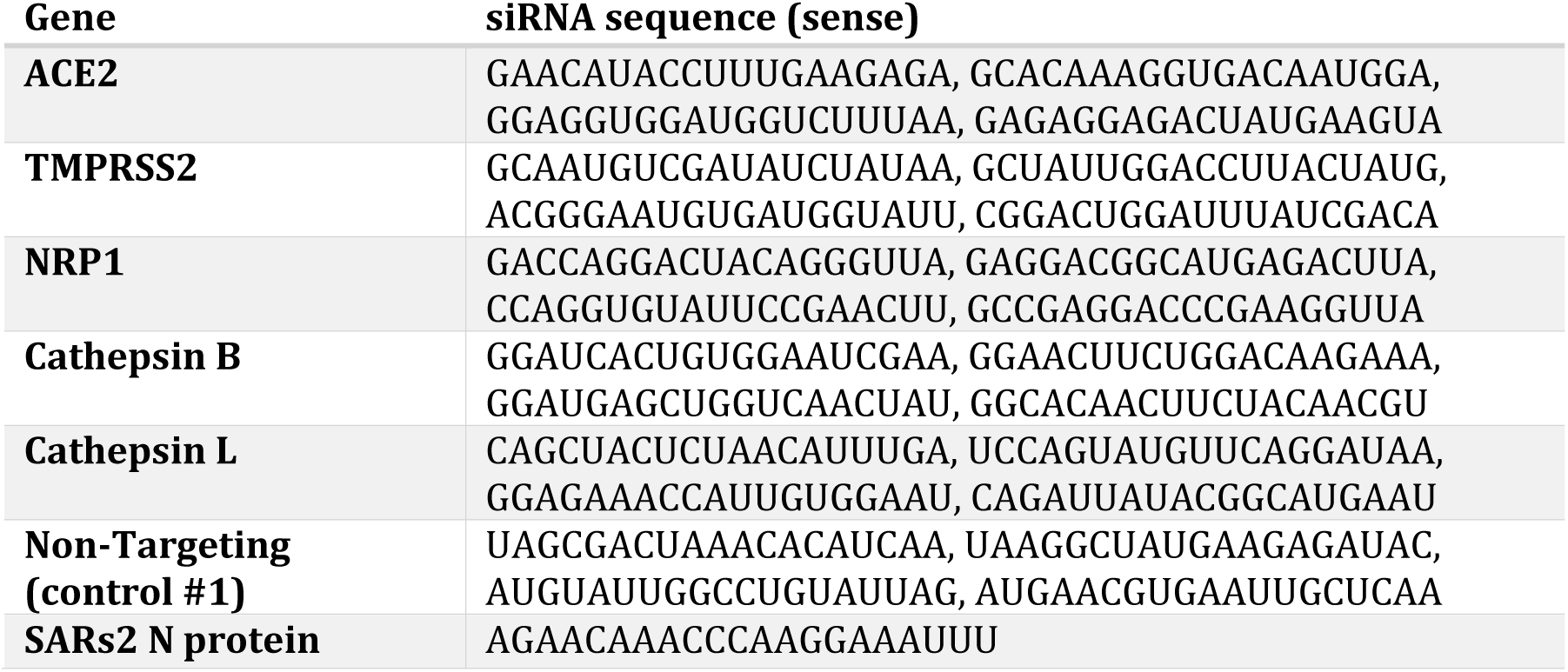
Dharmacon siRNA sequences used in this study.

### Cell Viability Assay

Caki-1 cells were seeded onto a black-sided, clear-bottomed 96-well plates after reverse transfection with siRNA (20 nM, as described above) and incubated for 72 h. CellTiter-Blue (Promega, G8080) reagent was added to each well at a 1:10 ratio. Following 3 hours of incubation at 37°C, fluorescence as a result of metabolic activity was measured using the BMG Clariostar plate reader (excitation: 545-20, emission: 600-40, 30 flashes/well, bottom optic) and fluorescence intensity was compared between the untreated and the drug treated cells to calculate a percentage viability.

### HEK-Blue Interferon Assay

Caki-1 cells were seeded in a 12-well plate and stimulated by either reverse transfecting with Poly I:C (Enzo) using Dharmafect 1 (working concentration of 0.1%), or with media containing Poly I:C (Enzo) or LPS (Sigma) at the indicated concentrations. At 24 hours post treatment, supernatant was harvested and 20 μl added to confluent HEK-Blue IFNα/β cells for 24 h. 20 μl of induced HEK-Blue cell supernatant was added to a corresponding plate with 180 μL of Quanti-Blue substrate in each well. After 3 h incubation SEAP levels were detected using the Clariostar (BMG) at 640 nm.

